# *Myc* is dispensable for cardiac development in the mouse but rescues *Mycn*-deficient hearts through functional replacement and cell competition

**DOI:** 10.1101/357335

**Authors:** Noelia Muñoz-Martín, Rocío Sierra, Thomas Schimmang, Cristina Villa del Campo, Miguel Torres

**Author notes:** Co-Corresponding authors: Cristina Villa del Campo and Miguel Torres Cardiovascular Development Program, Centro Nacional de Investigaciones Cardiovasculares, CNIC. Melchor Fernández Almagro 3, Madrid 28029, Spain.

## Abstract

Myc is considered an essential transcription factor for heart development, but cardiac defects have only been studied in global Myc loss of function models. Here, we eliminated Myc by recombining a *Myc* floxed allele with the *Nkx2.5Cre* driver. We observed no anatomical, cellular or functional alterations in either fetuses or adult cardiac Myc-deficient mice. We re-examined Myc expression during development and found no expression in developing cardiomyocytes. In contrast, we confirmed that *Mycn* is essential for cardiomyocyte proliferation and cardiogenesis. Mosaic Myc overexpression in a *Mycn*-deficient background, shows that Myc can replace Mycn function, recovering heart development. We further show that this recovery involves the elimination of Mycn-deficient cells by Cell Competition. Our results indicate that *Myc* is dispensable during cardiogenesis and adult heart homeostasis and *Mycn* is exclusively responsible for cardiomyocyte proliferation during heart development. Nonetheless, our results show that *Myc* can functionally replace *Mycn*. We also show that cardiomyocytes compete according to their overall Myc+Mycn levels and that Cell Competition eliminates flawed cardiomyocytes, suggesting its relevance as a quality control mechanism in cardiac development.

## Introduction

Myc transcription factors promote cell growth and division, being essential for proliferation in healthy tissues and tumours. Myc proteins belong to the bHLH-domain family and exert their functions mainly by regulating transcription. There are three members of the Myc family of transcription factors in mammals: Myc, Mycn and Mycl. All three transcripts show spatially-restricted patterns during postimplantation embryonic development. Deregulation of these genes has been linked with tumour formation and cell growth.

Myc expression is required for normal embryonic development in mammals, displaying widespread expression from early stages of development, and becoming regionally restricted starting at E7.5. Global Myc KO embryos die between days E9.5 and E10.5, showing defects in heart, pericardium, neural tube and delay or failure of embryo turning^1^. Strong Myc overexpression in transgenic mice enhances myocyte proliferation during heart development, promoting cardiac hyperplasia, which suggested the idea of an essential role of Myc in cardiomyocyte growth and proliferation during development^2,3^. In contrast, strong Myc overexpression during postnatal life leads to premature cardiomyocyte hypertrophy^3, 4^ and heart-specific deletion of Myc prevents hypertrophic growth in response to hemodynamic and pharmacologic stimuli^5^ and cold-induced hypertrophy^6^. Myc mRNA levels however, decrease in correlation with the transition from hyperplastic to hypertrophic growth^7^ and is not expressed in adult cardiomyocytes under normal conditions but becomes strongly activated following hypertrophic stimuli^8, 9^, which suggests that the physiological function of Myc in postnatal cardiomyocytes is restricted to the hypertrophic response to a challenge. In accordance to this idea, Myc deletion in cardiomyocytes of unchallenged adult mouse hearts does not lead to cardiac function alterations^5^.

Further experiments in a model of moderate overexpression of Myc produced a very different set of results. Mild Myc overexpression in a cellular mosaic fashion does not produce overt phenotypical alterations during embryonic development or adult life, but induces the phenomenon of Cell Competition, by which cells with enhanced anabolism eliminate and replace neighbours without altering tissue homeostasis^10, 11^. In cardiac-specific models of Myc mosaic overexpression at moderate levels, Myc-enhanced cardiomyocytes trigger the elimination of neighbouring wild type cardiomyocytes both during development and in the adult heart^12, 13^.

The changes induced by Myc moderate overexpression in cardiomyocytes remain within homeostatic limits both during development and in the adult heart^12^. Notably, in these experiments, Myc-enhanced adult hearts are not prone to hypertrophy but display a mild hyperplasic phenotype^12^. The contrast of these results with those obtained by strong overexpression of Myc in transgenic mice^3, 4^ suggests that the effects of Myc overexpression critically depend on the levels induced.

While the results obtained in overexpression experiments suggest a role for Myc during cardiomyocyte development, there are no studies reporting cardiac-specific deletion of *Myc*. Furthermore, the conditional deletion of *Myc* in the blood/endothelial lineage produces heart defects similar to those observed in the complete *Myc* elimination^14^ raising the possibility that the cardiac defects observed in the global mutant do not result from a primary function in cardiomyocytes. In contrast, *Mycn* is essential for cardiomyocyte development in conditional deletion models^15^. Myc and Mycn show high sequence and structure homology and this translates into a highly conserved function, as exemplified by Mycn full rescue of the Myc global knockout in a knock-in replacement mouse model^16^.

*Mycn*global mutants die in utero between E10.5 and E11.5, displaying smaller size and hypoplastic heart^17–20^, which is reproduced in a cardiomyocyte-specific deletion of *Mycn* using a *cTnT-Cre* driver^15^. Mycn is required for ventricular wall morphogenesis through its role in regulating compact layer cardiomyocyte growth, proliferation and maturation. The defects in heart growth were exclusively due to the reduction in proliferation and not to increased cell death^15^.

Here we studied the role of *Myc* during heart development, the ability of *Myc* to rescue *Mycn* deficiency during cardiogenesis and the involvement of Cell Competition and cardiomyocyte replacement in this rescue. We report the absence of Myc expression or function during cardiac development and the ability of *Myc*-expressing cardiomyocyte populations to repopulate *Mycn*-deficient hearts and to rescue Mycn function. Our results indicate that, while Mycn is an essential constitutive component of cardiomyocyte development, Myc is not involved in normal cardiomyocyte formation. Nonetheless, Myc is able to mimic Mycn function, promoting the elimination of Mycn-deficient cells to restore a viable heart.

## Methods

### Mouse Strains

*iMOS* Mouse lines have been previously described^10^. Homozygous *iMOS* females were mated with males carrying different Cre lines; *Wt1Cre*^21^, *Nkx2.5-Cre*^22^ to generate embryos. The *Mycn* floxed allele has been previously described^23^, as has been the Myc-GFP reporter^24^ the Mice were genotyped by PCR. All animal procedures were conducted in accordance with applicable institutional guidelines.

### ISH

WM ISH was performed on E9.5 embryos and E12.5 hearts as described previously, using Myc probe^10^

### Confocal microscopy

Histological sections and whole-mount embryos were imaged with a Nikon A1R confocal microscope using 405, 458, 488, 568 and 633 nm wavelenghts and 20x/0.75 dry and 40/1.30 oil objectives. Cardiomyocyte nuclei were counted using the cell counter tool of Image J. To estimate cardiomyocyte size, the number of nuclei was divided by the myocardial area calculated using the threshold detection of Image J. Areas occupied by EYFP and ECFP cells and EYFP and ECFP cell number were quantified using the threshold detection and particle analysis tools of the Image J, (NHI, http://rsb.info.nih.gov/ij). To calculate the relative frequency of ECFP cells, the percentage of ECFP cells in each embryo was divided by the average percentage in *iMOS*^*WT*^ mosaic. ECFP was scored by subtracting the information of EYFP from the anti-GFP staining which detects both fluorescent proteins.

### Adult measurements

After sacrifice, mice were weighed and hearts were extracted and rinsed in PBS. Hearts were weighed and tibia length of the posterior left leg was measured with a caliber.

### Immunofluorescence

Embryos were fixed overnight at 4ºC in 2% paraformaldehyde in PBS and whole-mount stained or embedded in gelatin and cryosectioned. Embryonic hearts were fixed in 2% PFA overnight at 4ºC and stained in whole-mount. Adult hearts were perfused and fixed in 2% paraformaldehyde in PBS 24 hours at 4ºC and paraffin-embedded for sectioning. Primary antibodies used were PCM1 (Sigma), α-SMA (Sigma), Myc (Millipore), Living colors Rabbit polyclonal anti GFP antibody (Clontech), c-TnT (Thermo Scientific). Immunofluorescence was performed following standard procedures. TUNEL was performed on heart sections using terminal deoxynucleotidyl transferase (TdT) and biotin-16-2-deoxyuridine-5-triphosphate (Biotin-16-dUTP) (both from Roche), and developed with 647-conjugates Streptavidin (Jackson ImmunoResearch). E9.5 embryos were cleared before whole mount confocal acquisition using ethyl cinnamate as described in^25^.

### Echocardiography study

Transthoracic echocardiography was performed blinded by an expert operator using a high-frequency ultrasound system (Vevo 2100, Visualsonics Inc., Canada) with a 40-MHz linear probe on a heating platform. Mice were lightly anesthetized with 0.5-2% isoflurane in oxygen, adjusting the isoflurane to maintain heart rate at 450±50 bpm. A base apex electrocardiogram was continuously monitored. Images were analysed using the Vevo 2100 Workstation software. Parasternal standard, 2D and MM, long and short axis view at the level of the papillary muscles (LAX and SAX view, respectively) were acquired.

### Statistical analysis

Expected versus observed frequencies were compared using the Chi square method.

Adult heart parameters were analyzed with unpaired T-test comparing Wt vs Myc-KO and Wt vs Het separately. Nuclei/myocardium area data was analyzed with Two-way ANOVA. To compare average percentages of ECFP cells between more than two groups, Kruskal-Wallis test was used (assuming non-normal distributions). For comparisons of two groups a Man-Whitney test was used. All comparisons were made using Prism statistical software.

## Results and discussion

### Myc is dispensable for heart development and adult heart homeostasis

To study Myc’s role during heart development, we conditionally deleted *Myc* using the *Nkx2.5-Cre* strain, which drives widespread Cre-mediated recombination in all cardiac precursors from around E8.0^22^. Embryos resulting from elimination of Myc function in cardiac progenitors (cKO-Myc) were viable and did not display any phenotypic abnormality (Figure 1A). cKO-Myc mice reached adulthood in the expected proportions and presented normal cardiac morphology (Fig 1B, C).

**Figure 1.**
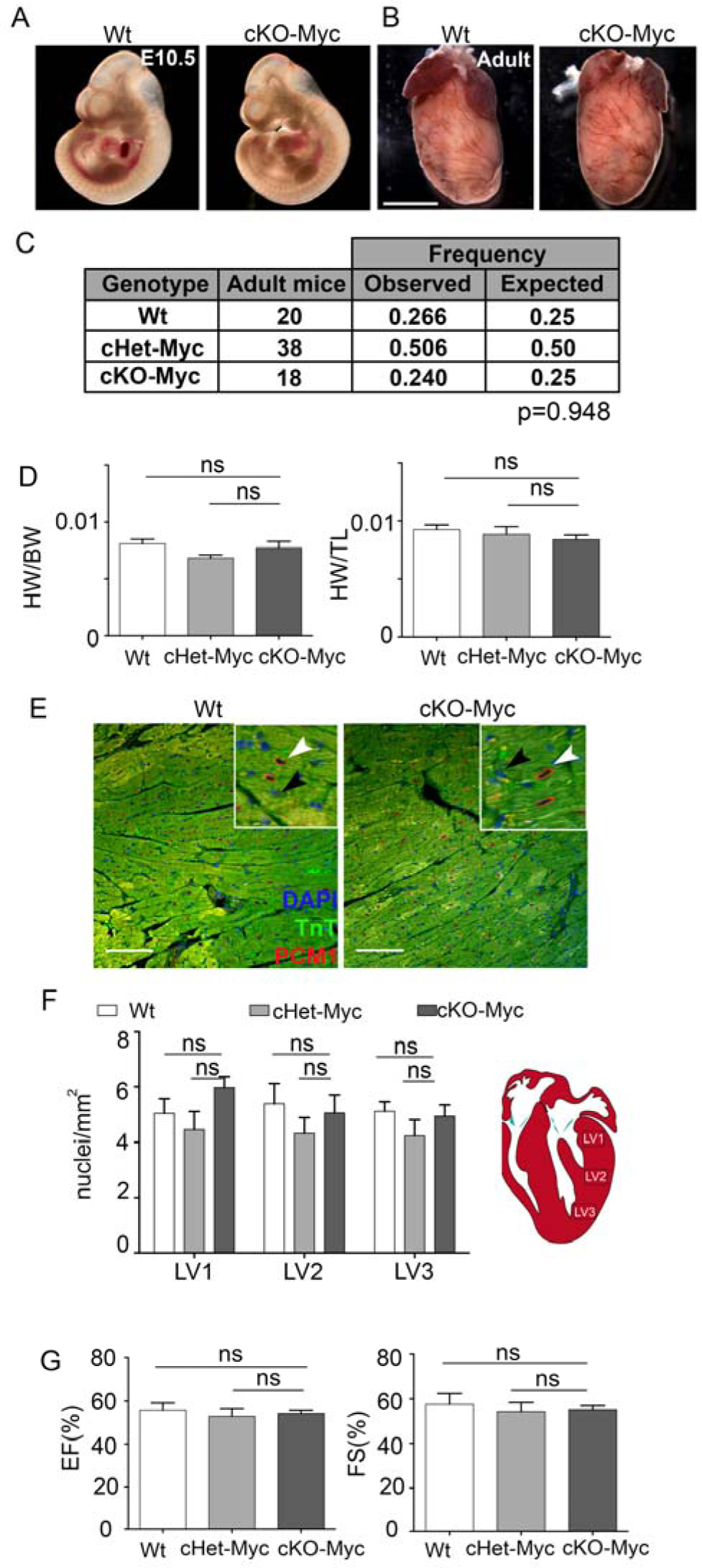
Myc is dispensable for heart development and adult heart in homeostasis. **A**. Whole mount wild type E10.5 embryo (left) and cKO-Myc E10.5 embryo (right). **B**. Whole mount wild type adult heart (left) and cKO-Myc adult heart (right). **C.** Table depicting the observed and expected frequencies of adult mice from the different genotypes. **D.** Graphs depicting the heart/body weight and heart/tibia length ratios in 10-week-old animals. **E.** Confocal images from sections of adult hearts stained with anti-PCM1 (red) and anti-TnT (green). **F.** Graph showing the quantification of CM nuclei per area in three different regions of the left ventricle. Location of the regions within the left ventricle is identified in the schematic as LV1, LV2 and LV3 Bar: 100um Data in **D, F, G** are means ±SEM; ns: p>0,05. N=3-8 mice/condition

Measurements of heart weight revealed no significant differences in size between cKO-Myc homozygous, heterozygous and wild type hearts (Fig 1D). To assess whether Myc loss had any effect on cardiomyocyte size or number, we estimated the density of cardiomyocyte nuclei in histological sections from different regions of the left ventricle (LV) (Fig 1E). The density of cardiomyocyte nuclei was similar between cKO-Myc homozygous, heterozygous and control hearts (Fig 1F). These data, together with the consistency in heart size, suggests that neither cardiomyocyte size nor number are changed by the loss of Myc expression.

To functionally assess cKO-Myc hearts, we performed echocardiographic assays on 10-week adult mice. No significant differences were found between groups in ejection fraction and fractional shortening parameters, indicating that the function of cKO-Myc hearts is not affected by the loss of *Myc* (Fig 1G). Overall, cKO-Myc hearts display normal morphology and function and therefore our data indicate that Myc is dispensable for heart formation and adult heart homeostasis.

### Myc is not detectably expressed in developing cardiomyocytes

The results obtained could be explained by the lack of Myc function during cardiomyocyte development or by the compensation of a putative Myc function by Mycn. *Myc* RNA expression has been reported by Northern blot in mid-gestation samples from whole myocardium^2, 7^ and Myc protein expression has been reported by Western blot from whole adult myocardium^5^. Here, we performed *in situ* analyses to determine which cells express Myc during myocardial development. We first performed *Myc* mRNA *in situ* hybridization (ISH) detection during embryogenesis^10^. At embryonic day 9.5 (E9.5) *Myc* displayed expression in the neural tube, branchial arches, cephalic regions and other non-cardiac tissues (Fig 2A), which is in agreement with previous reports^26^. At this stage *Myc* mRNA was not detected in the heart tube, while within the cardiogenic region, expression was seen in the proepicardium (Fig 2A). Analysis at later stages showed weak Myc mRNA detection in the distal outflow tract (OFT) and subepicardium at E12.5 (Fig 2B). These results were contradictory with our previous characterization of Myc protein distribution using anti-Myc antibodies in immunofluorescence, in which a clear signal was detected in cardiomyocytes at E10.5^12^. To resolve this contradiction, we repeated the immunofluorescence comparing wild type and cKO-Myc hearts at E10.5 (Figure 2 C, D). Detection of Myc expression in wild type embryos clearly identified a nuclear signal in cardiomyocytes (Fig 2C). This signal remained unchanged in cKO-Myc hearts (Fig 2D). This result contrasts with the observation that this antibody has been validated for endogenous Myc detection in the E6.5 mouse epiblast^10^. These results suggest that Myc antibody cross-reacts with other members of the family in cardiomyocytes, most likely Mycn, whose role and expression in cardiomyocyte development is well established, while it is not expressed in the E6.5 epiblast^15, 18^.

**Figure 2.**
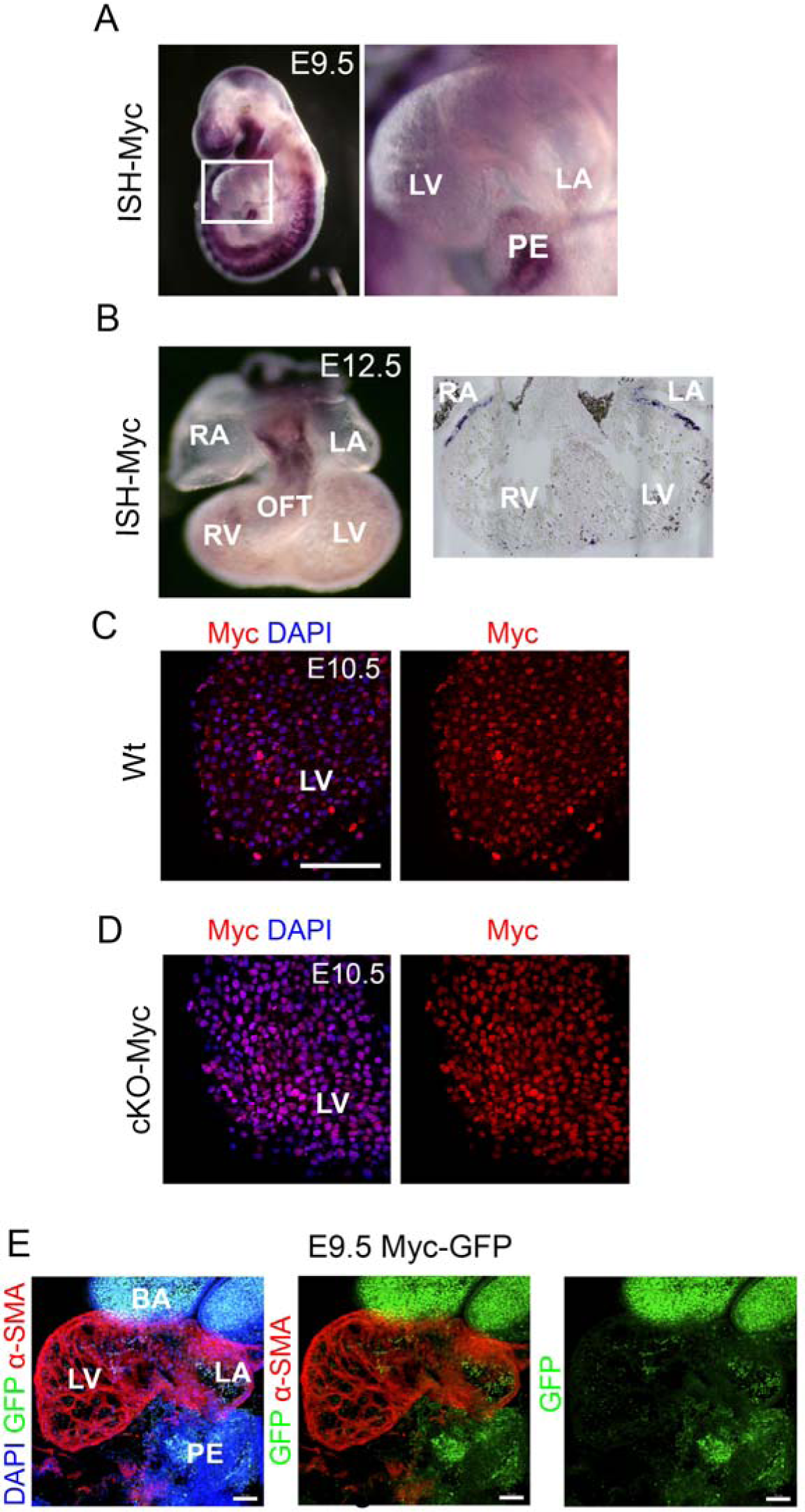
Myc is not widely expressed in cardiomyocytes during heart development. **A.** Whole mount in situ hybridization in E9.5 WT embryo for Myc, showing no expression in the heart (box in **A**). **B.** Whole mount in situ hybridization of a E12.5 WT embryonic heart for Myc. **C-D.** Confocal section of a whole E10.5 heart showing staining for *Myc* and DAPI (left) and *Myc* (right) for Wt **(C)** and cKO-Myc **(D)** embryo. **E.** Confocal section of a whole mount E9.5 *Myc-GFP* embryo showing GFP Myc expression and a-SMA. Bar, 50 μm in C, D and 70 in E. LV: Left ventricle, RV: Right ventricle, RA: Right atria, LA: Left atria, OFT: Outflow tract, BA: Branchial arches, PE: Proepicardium

To further confirm Myc expression in the developing heart, we took advantage of a *Myc-GFP* knock-in reporter line in which endogenous Myc protein expression is reported by the green fluorescent protein (GFP) fused to the endogenous Myc open reading frame^24^. In accordance to our ISH results, Myc-GFP expression at E9.5 was strongly detected in the branchial arches (Fig 2E) and in the proepicardium; however, no Myc expression was detected in the myocardium at this stage.

We conclude that Myc does not play a role in cardiomyocyte development because it is not expressed in this lineage and thus it does not act in redundancy with Mycn.

### Forced Myc expression in a mosaic fashion is sufficient to rescue *cKO-Mycn* cardiac defects

A relevant question is whether the different effects reported for *Myc* overexpression in cardiomyocytes result from Myc mimicking Mycn function by Myc. As mentioned above, *Mycn* can replace *Myc* functions when knocked-in to the *Myc* locus^16^. Here we addressed whether *Myc* could replace *Mycn* function in the developing heart. To test this, we used Myc overexpression from the Cre-inducible *Rosa26R*-*iMOS* mosaic system. The *iMOS^T1Myc^* allele allows the induction of mild overexpression of Myc in a cellular mosaic fashion^10^ (Fig 3A). In this mosaic model, 75% of recombined cells overexpress Myc and are reported by EYFP expression, while 25% do not overexpress Myc and are reported by the ECFP expression. We established crosses to obtain embryos simultaneously deleted for a conditional *Mycn* allele (*cKO-Mycn*) and activated for *iMOS^T1Myc^* with the *Nkx2.5-Cre* driver (Fig 3B). Embryos in which *Mycn* has been deleted but the *iMOS* mosaic is not activated are not viable past E10.5-E11.5 (Fig 3B), in accordance with the phenotype previously reported for Mycn deletion in cardiomyocytes^15^. Similarly, *iMOS^T1Myc^* activation on a wild type or *Mycn*-heterozygous background (iMOS-Myc) does not produce any phenotypic alteration^12^. Interestingly, cKO-Mycn littermates in which the *iMOS^t1Myc^* mosaic has been activated (cKO-Mycn; iMOS-Myc) were viable and indistinguishable from iMOS-Myc littermates (Fig. 3B). Histological analysis and study of the contribution of the cells recombined by *Nkx2.5-Cre* at E13.5 showed normal contribution of cardiac progenitors to the heart and no morphological alterations in either cKO-Mycn or Mycn-Wt hearts with iMOS-Myc overexpression (Fig. 3C). These results indicate that mosaic overexpression of one dose of Myc, driven by the endogenous *Rosa26* promoter, is enough to functionally replace the loss of *Mycn* expression during heart development.

**Figure 3.**
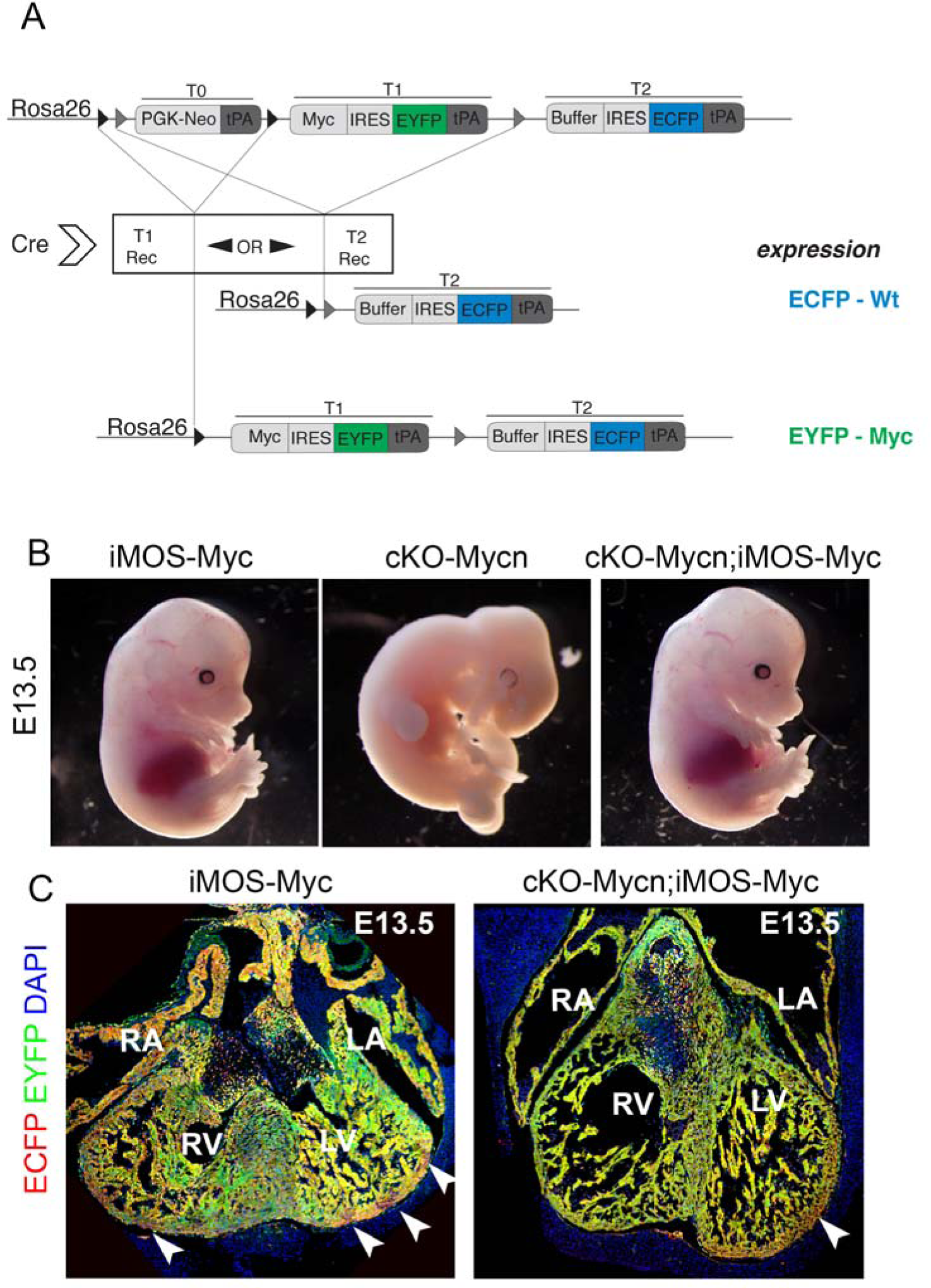
Myc expression in a mosaic fashion is sufficient to rescue Mycn embryonic lethal phenotype. **A.** Schematic of the *iMOS* construction. If T0 cassette is excised, T1 is expressed (EYFP-Myc). If T2 recombination takes place both T0 and T1 are excised leading to the expression of T2 (ECFP-Wt). This gives rise to two labeled cell populations at random under the control of Rosa26 promoter. **B.** Whole mount image of a E13.5 iMOS-Myc embryo (left),cKO-Myc embryo (middle) and cKO-Mycn; iMOS-Myc (right). **C.** Confocal section of a E13.5 heart from a iMOS-Myc embryo, showing EYFP-Myc and ECFP-wt cell populations (left) and cKO-Mycn; iMOS-Myc embryo, showing EYFP-Myc and ECFP-wt cell populations (right). Arrowheads point to ECFP positive cells. LV: Left ventricle, RV: Right ventricle, RA: Right atria, LA: Left atria. (N=3 cKO-Mycn; iMOS-Myc, N=3 cKO-Mycn, N=3 iMOS-Myc in Mycn heterozygous or wild type backgrounds).

### Cell Competition contributes to the rescue of *Mycn*-deficient hearts by stimulating the replacement of deficient cells

The complete phenotypic rescue of *cKO-Mycn* hearts suggested that, in addition to cell-autonomous replacement of Mycn function by Myc, some cell non-autonomous mechanism should operate to either eliminate or rescue the 25% of cells that do not activate *Myc*. To understand which of these mechanisms is at work, we determined the proportion of ECFP and EYFP cardiomyocyte populations in different genetic configurations. Control *iMOS* mosaics expressing only the fluorescent proteins over a wild type background produce a 25-75% distribution of ECFP and EYFP cardiomyocytes when activated by the Nkx2.5-Cre driver (iMOS-Wt) (Figure 4A,D) and the same experiment performed with the *iMOS^T1Myc^* mosaic reduces the cell population that does not overexpress Myc (ECFP cells) to 60% of the original, due to cell competition (Fig 4D)^12^. When the same experiment was performed over a cKO-Mycn heterozygous background, the proportion of ECFP cells observed was about 50% in E10.5 hearts (MycnHet; iMOS-Myc) (Fig 4B, D) whereas in a background homozygous for cKO-Nmyc (cKO-Mycn; iMOS-Myc), the proportion of ECFP cells dropped to 15% (Fig 4C, D). This elimination was further exacerbated in the cKO-Mycn homozygous background with progression of development, showing a proportion of only 4.5% of the original pool of ECFP cells at E13.5 (Fig 4F, D). These results suggest that not only the cell-autonomous replacement of Mycn by Myc, but also a replacement of *Mycn*-deficient cells by iMOS-Myc overexpressing cells contributes to the rescue of cKO-Mycn hearts.

**Figure 4:**
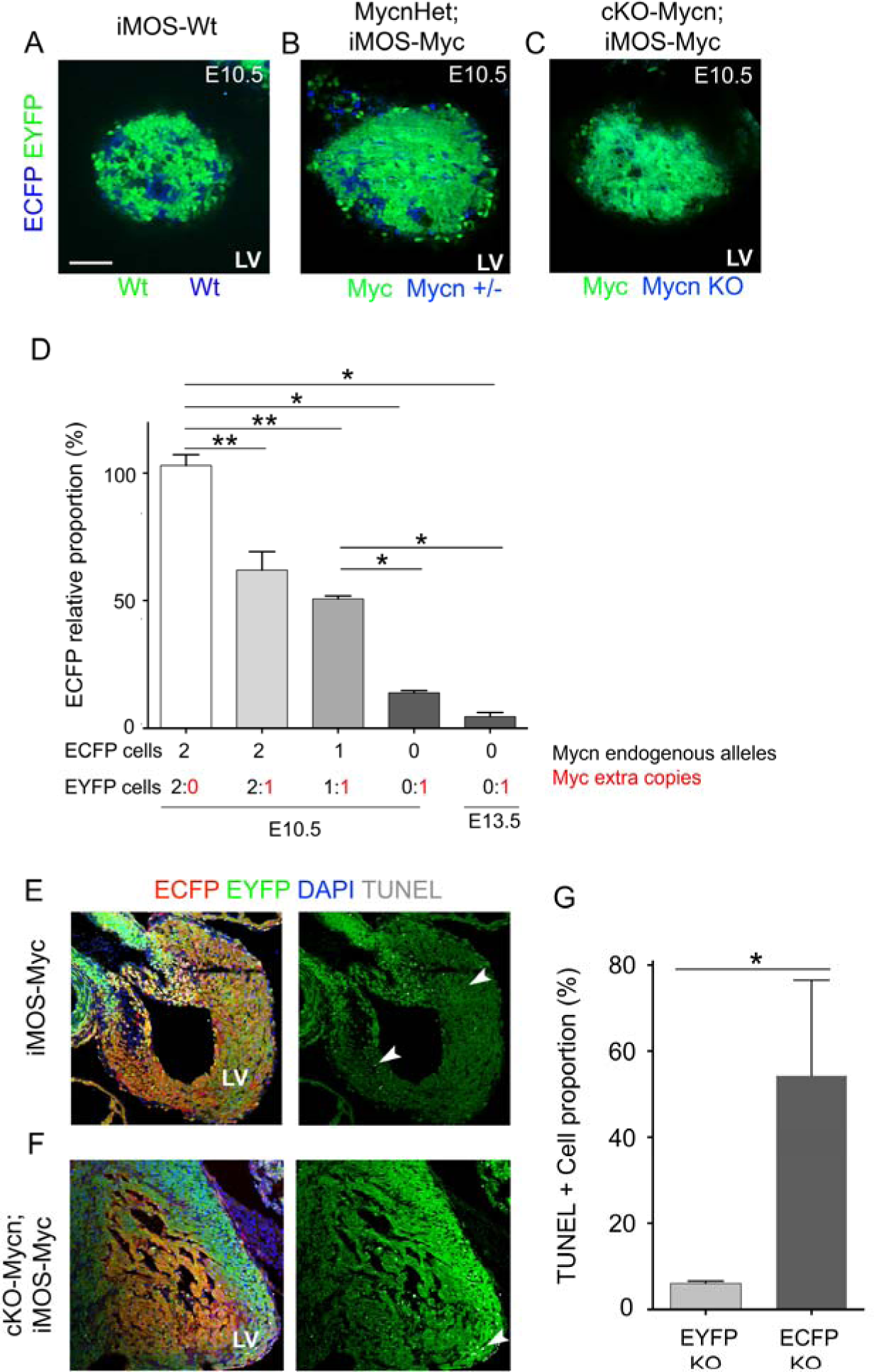
Myc and Mycn interplay can trigger a competitive process in cardiac progenitors, displacing those cells with lower relative Myc/Mycn levels, actively eliminated by cell competition. **A-C** Confocal sections from E10.5 hearts from iMOS-Wt (**A**), Mycn-Het; iMOS-Myc (**B**) and cKO-Mycn; iMOS-Myc (**C**) showing an overlay of EYFP and ECFP. **D.** Percentage of ECFP+ cells in hearts of the *iMOS-Myc* (MYC) mosaics in the three different Mycn backgrounds relative to that observed in the *iMOS-WT* (WT) mosaics, normalized to 100%. **E.** Confocal images of E13.5 heart sections of *iMOS-Myc* hearts showing ECFP and EYFP cell populations and TUNEL positive cells. **F.** Confocal images of E13.5 heart sections of cKO-Mycn; iMOS-Myc hearts showing ECFP and EYFP cell populations and TUNEL positive cells. **G**, Graph showing the relative percentage of TUNEL+ cells in EYFP and ECFP cell populations in cKO-Mycn;iMOS-Myc embryonic hearts. Bar 50 μm. LV: Left ventricle. Data in **D** are means ±SEM; * p<0,05; **p<0,01. Data in **G** are means ±SEM. Arrowheads in E and F point to TUNEL positive ECFP cells.

We next explored the possibility that the elimination of this cell population takes place by Cell Competition. Elimination of Mycn-deficient cells when confronted with Myc-overexpressing cells was much more efficient than elimination of wild type or Mycn-heterozygous cells, which would fit a scenario in which both Myc and Mycn act additively to determine cardiomyocyte competition ability. An alternative view would be that Mycn-deficient cells are not actively eliminated but just diluted out due to their limited ability to proliferate. To discriminate between these possibilities, given that Mycn-deficient cardiomyocytes do not autonomously display an increased apoptotic rate^15^ and that apoptosis is a hallmark of Myc-induced cell competition in the developing mouse heart, we performed TUNEL analysis.

As previously reported, in *iMOS^t1Myc^* mosaics induced over a wild type background the EYFP-Myc cell population shows nearly undetectable apoptosis, while the ECFP-WT cardiomyocyte population shows close to 6% apoptotic rate (Figure 4E)^12^. In contrast, in *iMOS^T1Myc^* mosaics induced over the cKO-Mycn background, while the apoptotic rate was also very low in EYFP-Myc cells, the apoptotic rate of the ECFP Mycn-KO cells increased over 50% (Figure 4F, G). These results indicate that confrontation with Myc-rescued cells produces a strong selective apoptotic elimination of Mycn-KO cells, otherwise viable in a homotypic environment^15^.

Taken together, our results show that Myc is not required for heart development and is not detectably expressed in developing cardiomyocytes. In the context of previous evidences from adult heart endogenous Myc expression and function analyses^2, 5, 7^, it is concluded that Myc expression and role in heart physiology is restricted to stress responses during adult life, while Mycn fully assumes the constitutive roles of the family during cardiogenesis. In addition, we show that Myc can replace Mycn in its functions during cardiomyocyte development and that Cell Competition contributes to rescuing heart function by stimulating the elimination of defective cells. This demonstration adds to previous evidence indicating the great plasticity of the developing heart to adapt to the progressive loss of up to 50% of its cardiomyocyte population by compensatory proliferation of the healthy population^27^. In contrast to this previously reported model, in which the unhealthy cardiomyocyte population is autonomously predisposed to cell death^27^, in the model presented here the unhealthy cell population is not prone to cell death when in isolation but undergoes massive cell death by confrontation with a Myc-rescued cell population. These results suggest an endogenous role for cell competition in the correction of contingent defects that may appear in cardiomyocytes during development.

## Acknowlegdments

We thank members of the Torres group for stimulating discussions and suggestions. We thank members of the microscopy and histopathology CNIC units, led by Valeria Caiolfa and Antonio de Molina-Iracheta, respectively, for excellent support and sample processing. We also thank the CNIC Advanced Imaging Unit and the CNIC Animal Facility personnel for their excellent work and support.

